# Wnt16 Elicits a Protective Effect Against Fractures and Supports Bone Repair in Zebrafish

**DOI:** 10.1101/2020.05.20.106328

**Authors:** Lucy M. McGowan, Erika Kague, Alistair Vorster, Elis Newham, Stephen Cross, Chrissy L. Hammond

## Abstract

Bone homeostasis is a dynamic, multicellular process which is required throughout life to maintain bone integrity, prevent fracture and respond to skeletal damage. *WNT16* has been linked to bone fragility and osteoporosis in humans, as well as functional haematopoiesis of leukocytes *in vivo*, but the mechanisms by which it promotes bone health and repair are not fully understood. We used CRISPR-Cas9 to generate mutant zebrafish lacking Wnt16 (*wnt16*^*-/-*^) to study its effect on bone dynamically. *wnt16* mutants displayed variable tissue mineral density and were susceptible to spontaneous fractures and the accumulation of bone calluses at an early age. Fractures were induced in the lepidotrichia of the caudal fins of *wnt16*^*-/-*^ and wild type (WT) zebrafish; this model was used to probe the mechanisms by which Wnt16 regulates skeletal and immune cell-dynamics *in vivo. wnt16* mutants repaired fractures more slowly compared to WT zebrafish. Osteoblast cell number was reduced at the fracture site 4 days post-injury in *wnt16* mutants, coinciding with prolonged activation of the canonical Wnt signalling pathway. Surprisingly, we found no evidence that the recruitment of innate immune cells to fractures was altered in *wnt16* mutants. This study highlights zebrafish as an emerging model for functionally validating osteoporosis-associated genes and investigating fracture repair dynamically *in vivo*. Using this model, we demonstrate that Wnt16 protects against fracture and is likely to support bone repair by attenuating the activation of the canonical Wnt signalling pathway to facilitate osteoblast recruitment and bone matrix deposition.

## Introduction

The maintenance of skeletal health is central to many essential processes in the body; in addition to facilitating movement and protecting vital organs, bones regulate mineral reserves, haematopoiesis and influence systemic hormone levels [1]. Skeletal homeostasis is maintained by numerous cell types such as chondrocytes, osteoblasts, osteocytes, osteoclasts and innate immune cells [2, 3]. These cell types act in concert to maintain an optimal balance between bone deposition and bone resorption under steady state conditions and respond to acute skeletal damage such as fracture [4, 5]. Osteoporosis occurs when bone deposition is reduced in relation to bone resorption, resulting in low bone mineral density (BMD) and loss of bone integrity [3]. Poor bone quality and low BMD is a strong predictor of fracture risk [6]. Currently, an estimated 3.5 million people in the UK suffer with osteoporosis, resulting in over half a million fractures per year [7]. Fragility fractures cause extensive morbidity and pose a high socioeconomic burden; as the ageing population increases, the treatment costs associated with osteoporotic bone fractures are set to rise by 30% in the next decade. Hence, there is an urgent unmet demand to understand the underlying causes of osteoporosis, identify novel targets for therapeutic intervention and promote optimal bone repair post-fracture.

Wnt signalling pathways are highly conserved, central regulators of skeletal development and homeostasis which act on bone throughout the lifetime of vertebrate organisms [8]. Canonical Wnt pathway activation in cells leads to the stabilisation of β-catenin and activation of transcription factors, whereas the calcium-dependent and planar cell polarity non-canonical Wnt signalling pathways regulate intracellular calcium levels and Jun N-terminal kinase (JNK) activity, respectively [9]. Wnt ligands are a family of secreted glycoproteins which influence cell stemness, proliferation, differentiation and migration via Wnt signalling pathways [10]. WNT16 is one such ligand which can influence the activity of canonical and non-canonical Wnt pathways [11, 12]. Recently, WNT16 has emerged as a potential regulator of cortical bone thickness and bone mineral density, with mutations in *WNT16* being linked to osteoporosis susceptibility in human genome wide association studies (GWAS) [13-15]. Furthermore, a meta-analysis of GWAS in women aged 20-45 years also associated *WNT16* with lumbar-spine BMD, indicating that *WNT16* may influence BMD throughout life, not only in post-menopausal populations [16].

Current experimental evidence highlights WNT16 as potential regulator of bone homeostasis and repair, as well as immune cell development. Knockout of *Wnt16* in mice has been shown to lead to decreased cortical bone thickness and up to a 61% decrease in femur and tibia bone strength compared to wild type littermates in three-point bending tests [17]. Whilst, loss of *Wnt16* in mice decreases bone strength, overexpression of *Wnt16* in osteoblasts (under the *Col1a1* promoter) leads to increased bone formation [18, 19]. However, one study showed that *Wnt16* overexpression in osteoblasts could not counter glucocorticoid-induced osteoporosis and bone loss, suggesting that other factors play a role [19]. One possible explanation could include interactions with the immune system. Glucocorticoid treatment in zebrafish has been demonstrated to suppress the innate immune system and osteoblast activity leading to decreased bone synthesis [20]. It has also been shown that morpholino-mediated knockdown of *wnt16* in zebrafish embryos results in impaired haematopoiesis and loss of thymic T lymphocytes at 4 days post-fertilisation (dpf) [21]. Embryonic knockdown experiments demonstrated that somatic *wnt16* expression is required for the upregulation of notch ligands and subsequent expression of the haematopoietic stem cell (HSC) marker *cd41*, which is needed for proper immune cell differentiation [21]. Despite its proposed role in early HSC development, the relationship between Wnt16 and the immune system has not been explored further in adult tissues or in stable mutant lines. Moreover, there is increasing interest in the interplay between immune cells and bone; osteoclasts and macrophages are derived from a common myeloid progenitor cell population and it is thought that macrophages can differentiate directly into osteoclasts in response to environmental molecular stimuli [22]. The rapid but tightly regulated recruitment of innate immune cells is also required for optimal bone repair post-fracture [23, 24]. WNT16 has been linked to bone maintenance, fracture susceptibility and leukocyte differentiation. However, functional studies to elucidate the role of WNT16 in these dynamic processes are still required.

Zebrafish (*Danio rerio*) serve as excellent models for studying both the musculoskeletal system and innate immunity. Approximately 85% of human disease-related genes have an ortholog in zebrafish [25]; as a result, many of the developmental processes, cell types and immune cell populations contributing to bone maintenance in humans are strongly conserved in zebrafish [26, 27]. Crucially, transparent zebrafish fin tissue provides optical clarity for high-quality, dynamic live imaging of adult bone tissue and injury repair *in vivo* [28]. Recently, the crushing of zebrafish caudal fin ray bones (lepidotrichia) was established as a model for studying fracture repair *in vivo* [29]. Therefore, we used CRISPR/Cas9 technology to generate a stable *wnt16*^*-/-*^ mutant line of zebrafish to investigate how loss of functional *wnt16* would affect bone maintenance, fracture repair and innate leukocyte function. We demonstrate that lack of Wnt16 in zebrafish leads to variable tissue mineral density in the fins and increased frequency of spontaneous fractures of caudal lepidotrichia in early adulthood. We employed an induced fracture model, to further characterise key immunological and osteological events underpinning bone repair in zebrafish. We show that *wnt16*^*-/-*^ zebrafish repair bone more slowly compared to wild type (WT) zebrafish. Impaired fracture healing in *wnt16*^*-/-*^ zebrafish coincided with higher levels of canonical Wnt activation and delayed osteoblast recruitment. Surprisingly, the recruitment of innate immune cells (neutrophils and macrophages) was unaffected by loss of Wnt16 post-fracture. We found no measurable difference in overall osteoclast activity (tartrate-resistant acid phosphatase (TRAP) staining) but observed more distinct, concentrated areas of TRAP labelling in *wnt16* mutant fractures. Taken together, our data suggests that Wnt16 promotes optimal bone repair post-fracture by regulating osteoblast activity and bone matrix synthesis via the regulation of canonical Wnt activity. This highlights the modulation of the canonical Wnt pathway and Wnt16 as potential osteo-anabolic candidates for further exploration in osteoporosis therapy development. Our data also further promotes zebrafish as a novel model for the dynamic study of fracture repair *in vivo* and rapid validation of human osteoporosis-associated genes.

## Materials and Methods

### Transgenic zebrafish lines and animal husbandry

All zebrafish were maintained at the University of Bristol’s Animal Scientific Unit as previously described [30, 31]. Experiments were approved by local ethical committee (the Animal Welfare and Ethical Review body for University of Bristol) and performed under a UK Home Office project license. Transgenics used have been previously described (Table 1).

**Table 1.** Transgenic lines as listed on zfin.org and abbreviations used in text.

| Line Name and Reference | Abbreviation | Description |
| --- | --- | --- |
| Tg(7xTCF-Xla:Siam:nlsGFP) [32] | Wnt:GFP | Canonical Wnt activity |
| *osx-nls:eGFP* [33] | *osx*:GFP | Osteoblasts |
| Tg(*col2a1aBAC:mCherry*) [34] | *col2a1*:mCherry | Chondrocytes |
| Tg(*mpeg1:mCherry*) [35] | *mpeg1*:mCherry | Macrophages |
| Tg(ET30:lyzC:DsRed) [36] | *lyzC*:DsRed | Neutrophils |

### wnt16 CRISPR mutant zebrafish

gRNAs were designed targeting exon 2 of *wnt16. g*RNAs were incubated with Cas9 protein (Thermo Fisher, B25641) prior to injections, performed at 1 cell stage eggs, as CRISPR/Cas9 mutagenesis was used to generate G0 mosaic zebrafish carrying indel mutations in exon 2 of *wnt16* as previously described in Brunt *et al*., [37]. G0s were raised to 3 months and crossed to wild type fish (TL/EKK strain) to generate heterozygous G1 embryos with a variety of *wnt16* mutant alleles. DNA was extracted from G1s, followed by PCR and cloning using TOPO-TA sequencing kit (Thermo Fisher), followed by sequencing. Two alleles were selected: *wnt16*^*bi667*^ (165 bp insertion, *wnt16*^*a1-/-*^) and *wnt16*^*bi451*^ (72 bp insertion, *wnt16*^*a2-/-*^), (Supplementary Figure S1 A). Both alleles lead to a premature stop codon compromising over 85% of the protein, likely resulting in nonsense mediated decay, therefore predicted to be null mutants (Supplementary Figure S1 B). Heterozygous *wnt16*^*+/-*^ fish were incrossed to generate stable homozygous (*wnt16*^*-/-*^) mutants which were used in experiments.

### Genotyping

Fish were genotyped by clipping the dorsal fin and placing the tissue in base solution (25 mM NaOH, 0.2 mM EDTA). Samples were heated to 98°C for 30 minutes and cooled to 4°C before neutralising with 40 mM Tris-HCl (pH 5.0). PCR was performed using EmeraldAmp^®^ GT PCR Master Mix and *wnt16* F-TTTTCCTCGGGCCTGGTTAT; R-GCCCTCTTTAACGCTCGGTA primers. Gel electrophoresis was performed using the PCR product from each sample (1.5% agarose in TAE + 1:10 000 SYBR Safe (Invitrogen)). Genotype was determined based on band separation due to variation in amplicon length: wild type = 216 bp, *wnt16*^*a1-/-*^ = 381 bp, *wnt16*^*a2-/-*^ = 288 bp (Supplementary Figure S1 C).

### Fracture induction

Young adult fish (6 months old) anaesthetised using MS222 (Sigma-Aldrich) and moved to a plastic dish for imaging. Lepidotrichia within the caudal fins were imaged prior to injury (see below). Fractures were induced by pressing on an individual segment of bone in the caudal fin lepidotrichia with a blunt-ended glass capillary tube. Fractures were induced proximal to the body of the fish, prior to the first bifurcation in the ray. Fish were recovered and reimaged at various time points post-injury.

### Imaging of fractures

Fish were housed individually and placed under anaesthetic at time points of interest post-fracture. Fractures were imaged in the dark using a DFC700T camera mounted to a MZ10F Stereomicroscope (Leica microsystems) before fish were revived immediately in fresh system water. Images were acquired using LAS X software 3.7.0 (Leica microsystems).

### Live staining of bone

To visualise bone repair in live zebrafish, various combinations of Alizarin Red stain (ARS) and calcein green staining were used. ARS was composed of 74 μM Alizarin Powder (Sigma-Aldrich) and 5 mM HEPES dissolved in Danieau’s solution [38]. Calcein green stain was composed of 40 μM calcein powder (Sigma-Aldrich) dissolved in Danieau’s solution (pH 8) [39]. Live fish were immersed in either ARS or calcein green for one hour, then immersed in fresh system water for 15 minutes prior to imaging to clear excess stain.

### Whole-mount fin immunohistochemistry

Whole fins were amputated and fixed in 4% paraformaldehyde overnight at 4°C. Fins were dehydrated in a series of increasing concentrations up to 100% methanol and stored at – 20°C. Fins were rehydrated and then washed 3 x in PBS-Tx (0.02% Triton-X in PBS) for 10 mins before 6ermeabilization in PBS-Tx + proteinase K (1:1000, Sigma Aldrich (P5568)) at 37°C for 90 mins. Solutions were refreshed every 30 minutes. Samples were washed 3 x in PBS-Tx for 10 mins and then blocked for 3 hours in blocking buffer (5% horse serum in PBS) and incubated in primary antibody overnight at 4°C. Samples were washed in PBS-Tx and blocked for 2 hours in blocking buffer staining with secondary antibody for 2 hours. Primary antibodies: pAb to WNT16 (abcam, ab189033, 1:300), mAb to GFP (abcam, ab13970, 1:500). The target human epitope of the polyclonal WNT16 primary antibody used was 50-60% conserved in zebrafish Wnt16, with predicted cross-reactivity at the C-terminus of the protein. Secondary antibodies: Alexa Fluor-647 and Alexa Fluor-488 (Thermo Fisher). Steps were performed at room temperature unless stated otherwise. Samples were mounted laterally in 1% agarose and imaged with a 10x objective lens on a SP5 confocal microscope (Leica Microsystems).

### Whole-mount larval immunohistochemistry

Larvae were euthanised in MS222 and fixed in 4% paraformaldehyde overnight at 4°C. Larvae were dehydrated in a series of increasing concentrations up to 100% methanol and stored at −20°C long-term until required. For staining, larvae were rehydrated and then washed 3 x in PBS-Tw (0.1% TWEEN-20 in PBS) for 10 mins. Larvae were permeabilised in PBS-Tw + proteinase K (1:1000) at 37°C for 25 mins (3 dpf) or 50 mins (5 dpf), with solutions refreshed after 30 mins. Samples were washed 3 x in PBS-Tw for 10 mins each and then blocked for 3 hours is blocking buffer before being stained and imaged as above. Larvae were mounted ventrally, and the jaw region imaged. Primary antibodies: chick α−L-plastin (gift from Martin lab [40]) and col2a1 (1:500, DSHB, M3F7). Secondary antibodies: Alexa-488, DyLight 550 (Thermo Fisher).

### Tartrate-resistant acid phosphatase (TRAP) staining

We used an Acid Phosphatase kit to detect osteoclast activity (Sigma-Aldrich, 387A). Fractures were induced in WT and *wnt16*^*-/-*^ *mpeg1*:mCherry zebrafish before being imaged and amputated at 0 hpi, 24 hpi, 4 dpi and 7 dpi. Amputated fins were stored on ice then fixed for 40 minutes at room temperature in TRAP-fix solution, comprised of 24% citrate solution (from kit), 65% acetone, 8% formaldehyde (37%) and 3% deionized water. Samples were washed in PBS-Tx 3 times. TRAP staining solution was prepared according to the kit instructions. Each fin was placed in a separate well of a 24-well plate and incubated at 37°C for 2 hours in 300 μl of TRAP stain. Fins were washed 3 times in PSB-Tx and post-fixed for 40 minutes at room temperature in 4% PFA before being transferred into 75% glycerol. Fins were stored at 4°C before imaging on a stereomicroscope.

### Micro computed tomography (μCT)

Adult fish were fixed in 4% PFA for 1 week followed by sequential dehydration to 70% ethanol. Fish were scanned using a Bruker SKYSCAN 1227micro-CT scanner with a voxel size of 5 µm, using an x-ray source of 60 keV, 50 W current and a 0.25mm thick aluminium filter. Each scan acquired 1500 angular projections with 400 ms exposure time over a 180° scan. X-radiographs were reconstructed using the filtered backprojection algorithm provided by NRecon software (v. 1.7.1.0) and saved as 8-bit tiff stacks. “Phantom” samples of known hydroxyapatite concentrations (0.25 and 0.75 g.cm^−3^ CaHA) were also scanned using identical settings to calibrate estimates of bone mineral density (BMD) in the micro-CT fin data.

### Tissue mineral density (TMD) analysis

Avizo image analysis software (version 8.0, Thermo Fisher Scientific) was used to generate 3D volume renders of whole fins using a combination of automatic and manual segmentation, which were saved as binary image stacks. The first 2 dorsal and ventral lepidotrichia were excluded from the analysis of all fins due to varying resolution. Image stacks were used to isolate the Greyscale values of segmented fins from values of surrounding soft tissue and air by multiplying these binary (fin = 1; non-fin = 0) stacks against the original reconstruction stacks using image algebra in Fiji/ImageJ [41]. Greyscale values within resulting stacks, where values > 0 consisted solely of those representing fins, were compared with the mean Greyscale values of both phantoms in order to calibrate the TMD values that they represent.

### Fluorescent image analysis

To quantify relative fluorescence intensities in fractures within transgenic fish, FIJI was used. The average intensity for each fracture within a region of interest (ROI) was measured and divided by the average intensity of uninjured bone in the same fish to give an “intensity ratio”; this analysis method normalises for variability of reporter expression between fish and allows for standardised comparison between individuals.

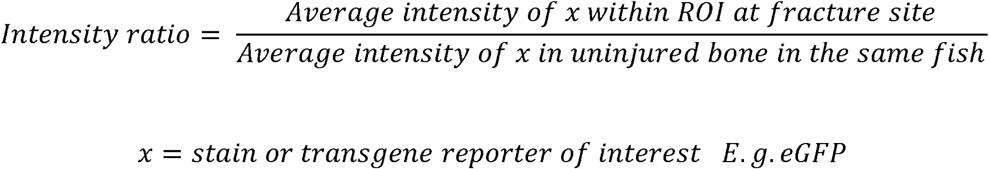

To analyse the number of immune cells responding to fracture, we used the freely available Modular Image Analysis (MIA; version 0.9.30) workflow automation plugin for Fiji [42-44]. Immune cell images were enhanced using the WEKA pixel classification plugin [45] and thresholded at a probability of 0.5. Adjacent cells in the binarised image were separated using an intensity-based watershed transform and individual cells subsequently identified as regions of connected foreground-labelled pixels [46]. Cells were subjected to a size filter, retaining only those in the range 30-500 μm^2^. The distance of each cell to the manually identified fracture site was measured.

### Statistical analysis

Statistical analyses were performed, and graphs were created in GraphPad PRISM 8 software. Where possible, a D’Agostino Pearson normality test was performed on data to determine whether a parametric or non-parametric statistical test should be used. Where two or more data sets were compared, a One-way analysis of variance (ANOVA) or a Kruskal-Wallis test was used to determine statistically significant differences between groups for parametric and non-parametric data, respectively. For comparison of WT and *wnt16* mutants throughout fracture repair, multiple t-tests were performed for each of the time points using the Holm-Sidak correction to calculate P values. Differences were considered statistically significant where P < 0.05.

## Results

### Young wnt16 mutant zebrafish are susceptible to spontaneous fractures which heal more slowly compared to wild type fish

*WNT16* has been associated with low eBMD and increased fracture risk [15, 17, 47], whilst *wnt16* mosaic mutant zebrafish displayed a low bone-mass phenotype [48]. Therefore, we used μCT to observe bone morphology and tissue mineral density (TMD) in whole-fins of adult WT and *wnt16*^*-/-*^ zebrafish. *Wnt16* mutants displayed a high degree of variability in TMD relative to WT specimens, as well as lower TMD (Figure 1 A-B). Images of *wnt16*^*-/-*^ fins showed a high frequency of bone calluses (Figure 1A), which form post-fracture and do not completely resolve after the bone has repaired [29]. Bone calluses in the caudal fin rays can be easily visualised using alizarin Red S (ARS). Thus, we next used ARS to compare the frequency of spontaneous lepidotrichia fractures in young, 6 month old (mo) WT and 6 mo *wnt16*^*-/-*^ uninjured fish. Bone calluses and spontaneous fractures were rarely observed in the 6 mo WT fish, with only 25% of fish sampled displaying a minimal number of calluses (≤ 3) (Figure 1 C-E). However, a significantly higher number of calluses were recorded in 6 mo *wnt16*^*-/-*^ fins; 100% of *wnt16*^*-/-*^ fins sampled contained calluses, with a mean of 8.5 calluses per fin versus 0.4 calluses per fin in WT. To test whether callus quantity increases with age, we quantified callus number in 20 mo and 30 mo WT fish. Aged WT fish were comparable in appearance and callus frequency to 6 mo *wnt16*^*-/-*^ fish (Figure 1 C-E). Collectively, this demonstrates that *wnt16*^*-/-*^ fish display a bone fragility phenotype predisposing them to spontaneous fractures and the accumulation of calluses at a young age.

**Figure 1.**
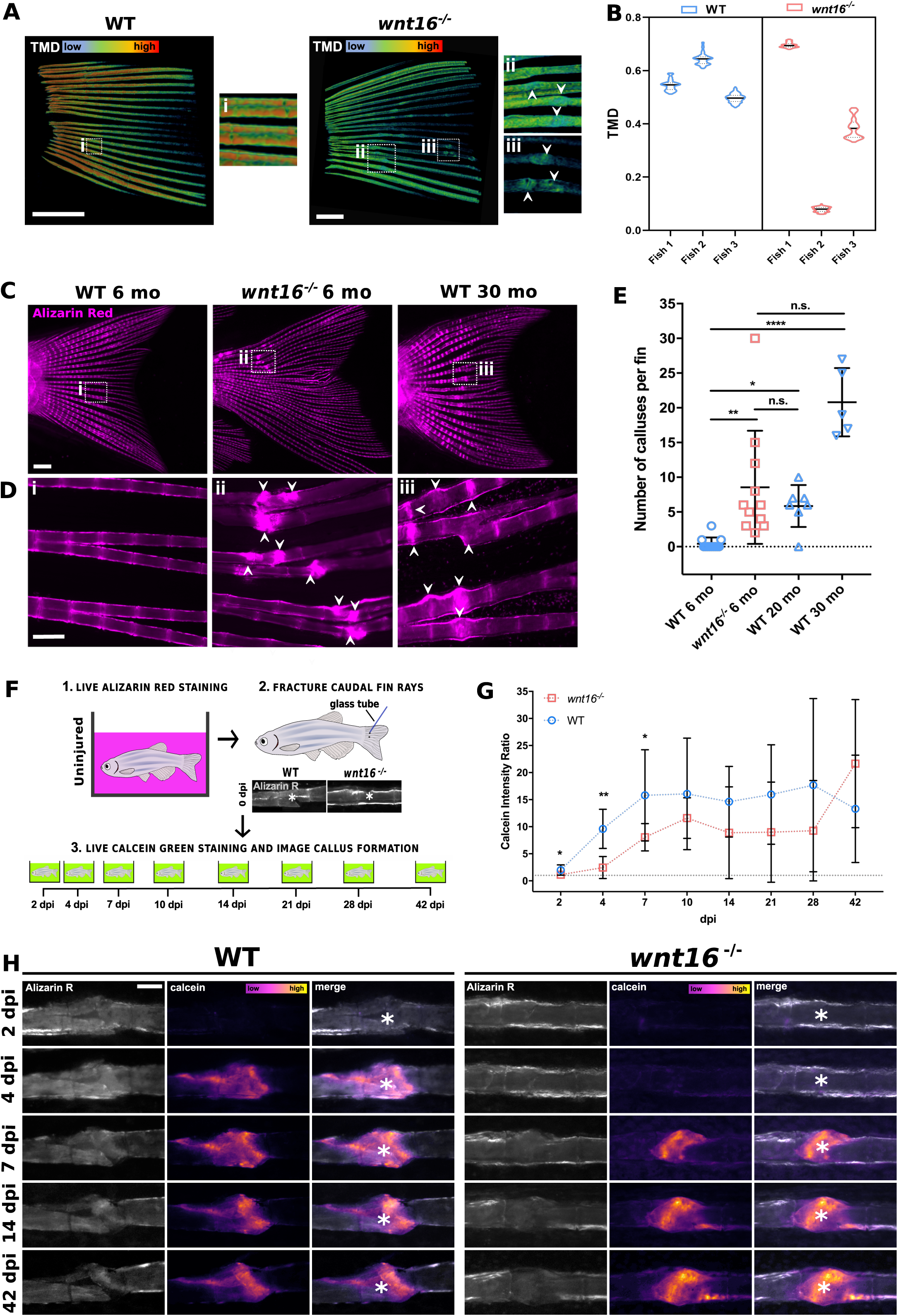
*wnt16* mutants are susceptible to lepidotrichia bone fractures which heal more slowly compared to wild type zebrafish. **A:** micro-CT images indicate lower and more variable tissue mineral density (TMD) and the presence of bone calluses (arrowheads) in the fins of *wnt16*^*-/-*^ zebrafish. **B:** Violin plots show distribution around mean (black line) TMD in wild type (WT) and *wnt16*^*-/-*^ fins. n= 3, scale bar = 1 mm. **C:** Uninjured WT and *wnt16*^*-/-*^ zebrafish were live-stained with alizarin red at 6, 20 or 30 months old (mo). Scale = 1 mm. **D:** Higher magnification of fins (from C) shows the presence of bone calluses (arrowheads) resulting from bone repair in 6 mo *wnt16*^*-/-*^ and 30 mo WT zebrafish but not 6 mo zebrafish. Scale = 200 μm. **E:** Quantification of bone calluses per fin shows that young *wnt16* mutants display a significantly higher number of calluses compared to WT fish at the same age but no significant difference compared with aged WT zebrafish. n ≥ 5 per condition. **F:** Schematic illustrating fracture induction assay and labelling of old bone (Alizarin R) and new bone (calcein green). **G:** Callus formation was quantified by measuring the calcein intensity ratio between the fracture-site and uninjured bone. Callus formation was significantly reduced from 2-7 days post-injury (dpi) in *wnt16* mutant compared to WT fractures. n ≥ 5 per condition. Grey dotted line indicates where calcein intensity at the fracture site = uninjured bone **H:** Representative images of WT and *wnt16*^*-/-*^ fish at selected time points post-injury show old bone labelled by Alizarin R (grey) and callus formation labelled by calcein (magenta = low intensity, yellow = high intensity). White asterisk = centre of fracture. Scale 200 μm. n.s = no significant difference, * = P < 0.05, ** = P < 0.01, **** = P < 0.0001.

### New bone matrix is incorporated more slowly post-fracture in wnt16 mutant zebrafish

Bone calluses are formed post-fracture; since *wnt16* mutants displayed variable TMD and a high number of calluses, we next tested whether fracture repair was impaired in *wnt16*^*-/-*^ zebrafish. Adult WT and *wnt16*^*-/-*^ fish were live stained in ARS to label bone and imaged prior to the induction of a fracture on a bone segment within the caudal lepidotrichia. Zebrafish were then live stained in calcein green to label newly incorporated bone matrix at the fracture site which was re-imaged at the time points indicated (Figure 1F). Injured *wnt16*^*-/-*^ zebrafish displayed significantly reduced callus formation within the first 7 days of fracture healing compared to WT, which was most apparent at 4 days post-injury (dpi) (Figure 1G and H). We also investigated whether chondrocytes were involved in lepidotrichia bone repair and if so, whether chondrocyte activity varied between WT and *wnt16* mutants. Fractures were induced in the caudal fins of transgenic *col2a1*:mCherry zebrafish (Table 1) and chondrocyte activity measured via fluorescence intensity. mCherry expression was almost undetectable throughout fracture repair and intensity ratios showed little variation from uninjured control bone at all time-points post-injury (Supplementary Figure S2 A-B). Moreover, no significant differences in Col2a1 levels were observed between WT and *wnt16* mutant fractures at any time point. This suggests that lepidotrichia fracture repair occurs predominantly via an intramembranous route and that chondrocytes are not required for adult bone repair in zebrafish.

### Osteoblast recruitment is delayed in wnt16^−/−^ zebrafish post-fracture

Osteoblast activation is a key event in the bone repair process post-fracture. Osteoblasts differentiate from mesenchymal stem cell (MSC) precursors expressing the transcription factor osterix (*osx*) and synthesise bone matrix within the initial soft callus; the callus hardens as it mineralises and is remodelled to restore the bone to a healthy state [49]. Moreover, transcriptomic analysis of osteoblast-prone clones isolated from tonsil derived MSCs showed that upregulation of WNT16 is predictive of osteogenic differentiation [50]. In zebrafish, osteoblasts dedifferentiate and proliferate in response to bone injury, migrating to the damaged tissue where they initiate bone repair [28]. Thus, we next investigated whether osteoblast number and distribution were impaired post-fracture repair in *wnt16*^*-/-*^ zebrafish. We performed live ARS prior to fin fractures of WT and *wnt16*^*-/-*^ zebrafish carrying the osteoblast-labelling transgene, *osx*:GFP (Table 1). Fractures were induced and re-stained with live ARS at the time points indicated to ensure labelling of any new bone prior to imaging (Figure 2A). The intensity of *osx*:GFP signal was measured as a ratio between the fracture site and uninjured bone and used as a proxy for relative increases in osteoblast number throughout fracture repair. In WT zebrafish, the number of osteoblasts at the fracture site peaked rapidly at 4 dpi, before steadily decreasing (Figure 2B & C). However, osteoblast number was significantly reduced at 4 dpi in *wnt16* mutants, not peaking until 10 dpi (Figure 2B & C). A comparable bony callus had formed at the fracture-site in both WT and *wnt16*^*-/-*^ by 15 dpi (Figure 2D). The delayed recruitment and reduced number of osteoblasts at 4 dpi coincided with slower callus formation in *wnt16*^*-/-*^ fractures (Figure 1 G-H). This demonstrates that osteoblasts in *wnt16*^*-/-*^ zebrafish can respond to bone injury but that the recruitment and activity of these osteoblasts are significantly delayed.

**Figure 2.**
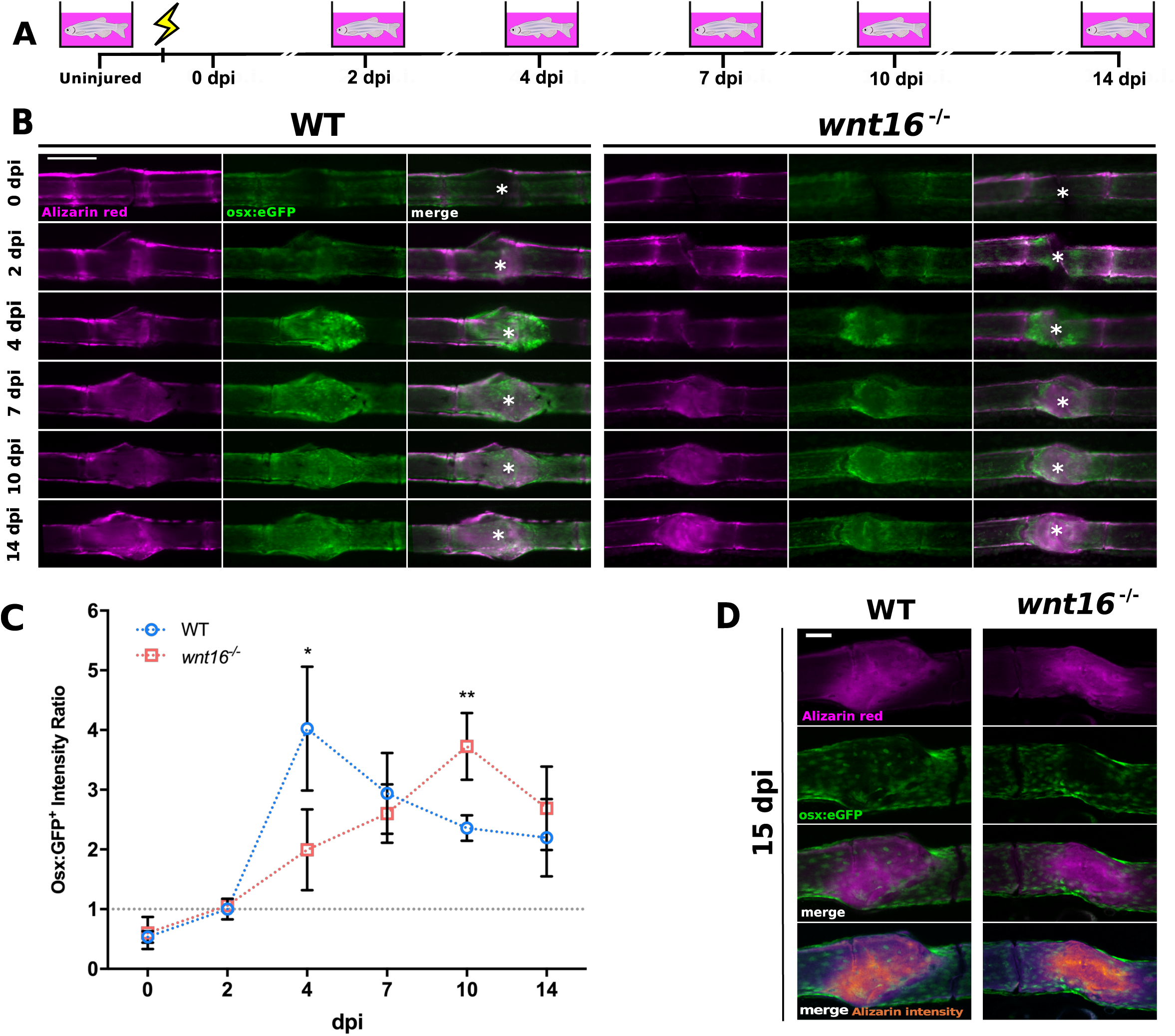
Osteoblast recruitment is significantly delayed post-fracture in *wnt16*^*-/-*^ zebrafish. **A:** Schematic illustrating induced-fracture time course and days post-injury (dpi) in which *osx*:GFP zebrafish immersed in Alizarin Red (magenta) for bone labelling and imaged. **B:** Representative images of calcified bone (Alizarin red) and osteoblasts (*osx*:GFP) at fracture site in wild type (WT) and *wnt16*^*-/-*^ throughout fracture repair. White asterisk = centre of fracture. Scale bar = 200 μm. **C:** Osteoblast number was quantified by measuring the fluorescence intensity of *osx*:GFP within the fracture site normalised to control bone in the same fin (intensity ratio). Grey dotted line indicates where osteoblast number at the fracture site = uninjured bone. Osteoblast recruitment was delayed in *wnt16*^*-/-*^ mutants, which had significantly fewer osteoblasts at the fracture site 4 dpi, but significantly more osteoblasts at 10 dpi compared to WT zebrafish. * = P < 0.05, ** = P < 0.01. n = 6 per genotype. **D:** Confocal imaging of bone in amputated fins at the end of the time course (15 dpi) shows complete union of fractures in both WT and *wnt16*^*-/-*^ zebrafish. Scale bar = 100 μm.

### Activation of the canonical Wnt signalling pathway is enhanced at the fracture site in wnt16^−/−^ zebrafish post-injury

Wnt signalling proteins regulate the stemness, differentiation and proliferation of MSCs and osteoblasts. Moreover, previous studies in mice have indicated that WNT16 may buffer levels of canonical Wnt signalling in response to injury [12]. Therefore, we investigated levels of canonical Wnt activity in *wnt16*^*-/-*^ zebrafish post-fracture using a β-catenin-responsive transgenic line (Wnt:GFP (Table 1)). Fractures were induced in the caudal lepidotrichia of the fish and imaged at identical time points as in Figure 2 A. In *wnt16*^*-/-*^ zebrafish, we observed a significant increase in the area of canonical Wnt-responsive cells at the fracture site from 2 dpi compared to WT (Figure 3). Canonical Wnt signalling remained elevated in *wnt16*^*-/-*^ fractures through to 4 dpi, where Wnt:GFP intensity ratios were comparable to WT fractures, before gradually decreasing to homeostatic levels by 10 dpi (Figure 3B). Fractured fins from WT and *wnt16*^*-/-*^ Wnt:GFP^+^ zebrafish were amputated and fixed 2 dpi and 4 dpi for immunohistochemistry to detect Wnt16, when the canonical Wnt pathway was most active. Immunohistochemistry revealed high levels of Wnt16 at the fracture site in WT zebrafish 2 dpi, coinciding with low levels of canonical Wnt pathway activation (Supplementary Figure S3 A). This was proceeded by a loss of Wnt16 signal at 4dpi, where canonical Wnt activation increased. Conversely, *wnt16*^*-/-*^ mutants displayed high levels of Wnt:GFP at both 4 and 7 dpi; only background levels of anti-Wnt16 immunolabelling were observed, which may be accounted for by antibody cross reactivity with other proteins (Supplementary Figure S3 B). Collectively, this suggests that enhanced canonical Wnt signalling underpins delayed callus formation and osteoblast differentiation in response to fracture in *wnt16*^*-/-*^ zebrafish.

**Figure 3.**
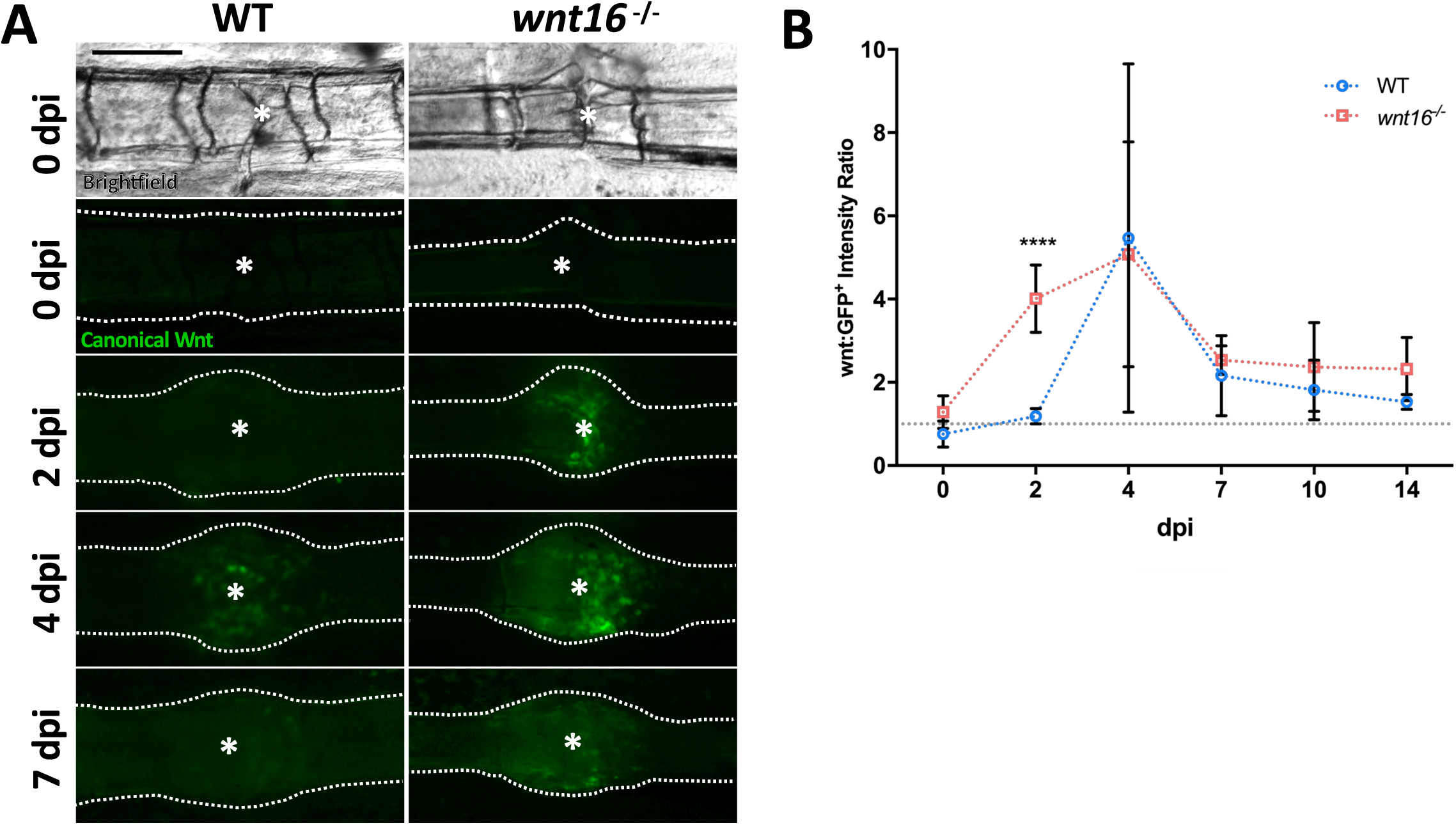
The Canonical Wnt pathway is over-activated in the early stages of fracture repair in *wnt16*^*-/-*^ zebrafish. **A:** Fractures were induced in *wnt*:GFP transgenic zebrafish which express GFP in cells responding to activation of the canonical Wnt signalling pathway. Representative images are shown from 0, 2, 4 and 7 days post-fracture (dpi). Dotted line = bone outline, White asterisk = centre of fracture. Scale bar = 200 μm. **B:** Levels of Wnt pathway activation throughout fracture repair were quantified by measuring the fluorescence intensity of Wnt:GFP within the fracture site normalised to control bone in the same fin (intensity ratio). Grey dotted line indicates where canonical Wnt activity at the fracture site = uninjured bone. *wnt16*^*-/-*^ zebrafish displayed significantly higher levels of canonical Wnt activity at 2 dpi compared to WT fractures. High levels of Wnt:GFP at the fracture site were sustained through to 4 dpi in *wnt16* mutants where they became comparable with WT. **** = P < 0.0001. n ≥ 6 per genotype.

### Innate immune cell dynamics are unaltered in wnt16^−/−^ zebrafish post-fracture

Fracture repair has been shown to comprise an inflammatory phase, a repair phase and a remodelling phase in mammals [51]. The controlled recruitment, activity and reverse migration of leukocytes during the inflammatory phase are known to be prerequisites for initiating osteoblast activity and optimal bone repair [51, 52]. Neutrophils are known to be amongst the first cells to be recruited to fractures to combat microbial infections and initiate bone repair [24]; stimulation of non-canonical Wnt pathways with recombinant WNT5a has been shown to initiate chemotactic migration and chemokine production in neutrophils, but whether WNT16 influences neutrophil recruitment is unknown [53]. Macrophages also rapidly respond to bone damage and continue to aid throughout the repair and remodelling phases in mammalian models of fracture [23]. A previous study indicated that *wnt16* expression was required for functional haematopoiesis in zebrafish embryos [21]. Additionally, overexpression of *WNT16* in mouse osteoblast-progenitor cells has been shown to partially rescue glucocorticoid-induced osteoporosis [54], suggesting that Wnt16 may regulate osteoblast activity and bone repair via immune cells. To validate whether early leukocyte development was impaired in *wnt16* mutants, we fixed zebrafish larvae at 3 and 5 days post-fertilization (dpf). Whole-mount immunohistochemistry was used to label cartilage in the developing skeleton (Col2a1) and immune cells (L-plastin) but surprisingly, no differences in leukocyte numbers were observed at either age (Supplementary Figure S4). Despite this, since callus formation and osteoblast differentiation were delayed in *wnt16*^*-/-*^ fractures, we also investigated whether immune cell recruitment to bone injury was altered in adult *wnt16* mutants. To this end, we used the *lyzC*:DsRed (neutrophils) and *mpeg1*:mCherry (macrophages) transgenic zebrafish lines (Table 1) to study leukocyte dynamics post-fracture in WT and *wnt16*^*-/-*^ zebrafish. Immune cell recruitment relative to the fracture site over time was quantified using modular image analysis [42]. The number of neutrophils (*lyzC*^+^ cells) and macrophages (*mpeg1*^+^ cells) within a 100 μm radius and 300 μm radius of the fracture were calculated (Figure 4 A). In both WT and *wnt16*^*-/-*^ fish, neutrophils were rapidly recruited to the fracture, peaking between 8 and 24 hpi (Figure 4 B). No significant differences in the number of neutrophils recruited to the fracture sites of WT and *wnt16*^*-/-*^ zebrafish were detected at any time point post-injury (Figure 4 C-D). Macrophages were also rapidly recruited to fractures in the first 24 hpi (Figure 4 E). Interestingly, we observed that macrophages responded to fracture in a biphasic manner, decreasing in number from 2-4 dpi, before peaking in number for a second time around 7 dpi (Figure 4 F-G). This suggests that phenotypically distinct populations of macrophages may be required at different stages post-fracture to contribute to efficient bone repair. Comparison between WT and *wnt16*^*-/-*^ zebrafish showed no difference in the number of *mpeg1*^+^ cells recruited to the fracture throughout repair, aside from a significant increase in macrophage number in *wnt16*^*-/-*^ zebrafish at 8hpi (Figure 4 F-G). These data suggest that, overall, leukocyte recruitment to fractures is not impaired in *wnt16* mutants and does not contribute to delayed bone repair resulting from loss of Wnt16.

**Figure 4.**
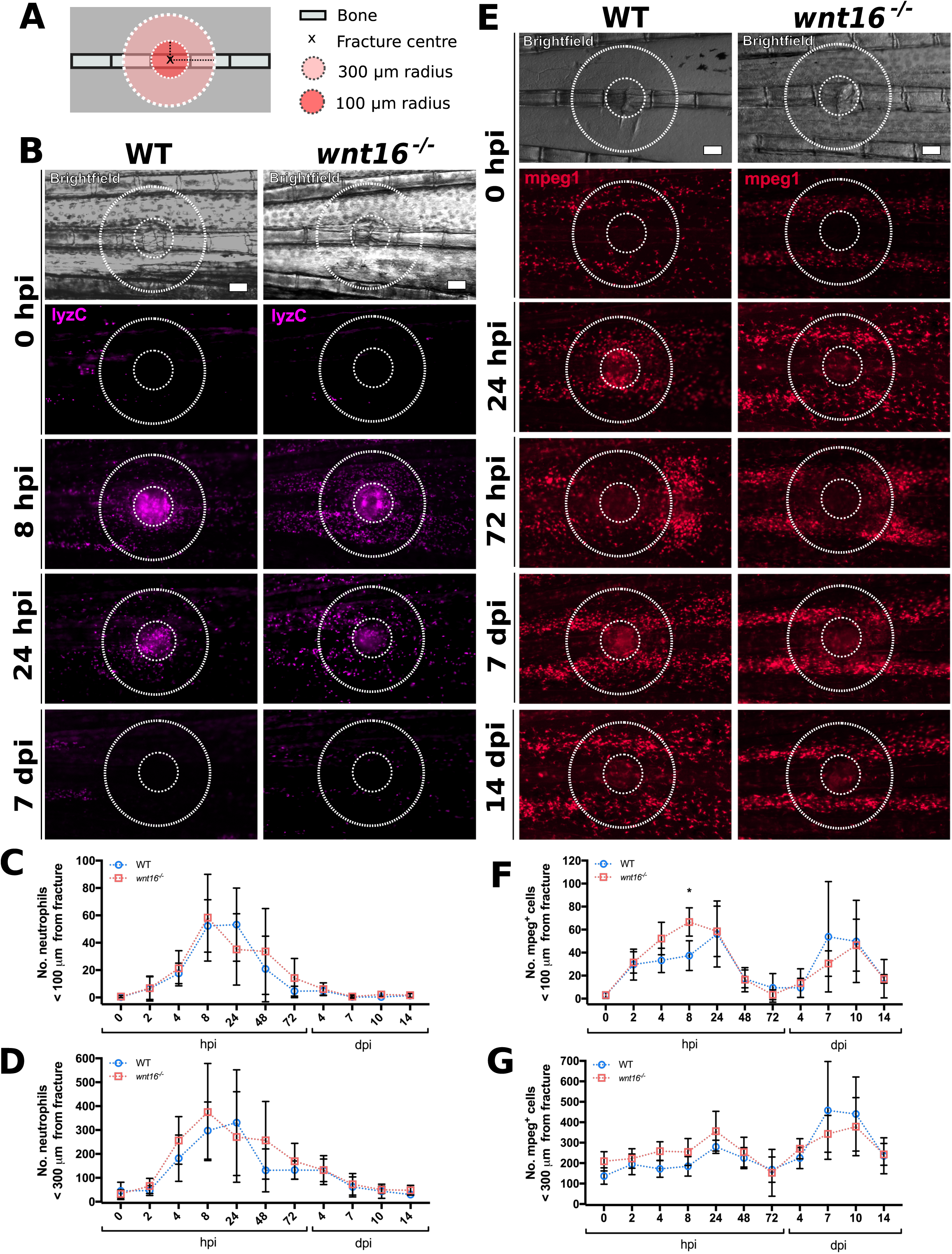
Loss of Wnt16 does not perturb leukocyte recruitment to bone post-fracture. Fractures were induced in wild type (WT) and *wnt16*^*-/-*^ zebrafish carrying *lyzC*:DsRed and *mpeg1*:mCherry transgenes to measure the recruitment of neutrophils and macrophages, respectively. **A:** Schematic depicting regions of interest around the fracture site where leukocyte recruitment was quantified. **B:** Representative images of from WT and *wnt16*^*-/-*^ zebrafish show neutrophil (*lyzC*^+^ cells) recruitment to fractured bone at 0, 8 and 24 hours post-injury (hpi) and 7 days post-injury (dpi). Scale bar = 100 μm. **C & D:** The number of neutrophils within 100 μm (C) and 300 μm (D) of the fractures were quantified in an automated manner using modular image analysis (MIA) from 0 hpi to 14 dpi. WT and wnt16 mutants displayed comparable numbers of neutrophils at the fracture site at all time-points post-injury. n ≥ 5 per genotype **E:** Representative images of from WT and *wnt16*^*-/-*^ zebrafish show macrophage (mpeg1^+^ cells) recruitment to fractured bone at selected time points from 0-14 dpi. Scale bar = 100 μm. **F & G:** The number of neutrophils within 100 μm (F) and 300 μm (G) of the fractures were quantified using MIA from 0 hpi to 14 dpi. WT and wnt16 mutants displayed comparable numbers of macrophages at all time-points post-injury, with the exception of 8 hpi when *wnt16* mutants had recruited significantly more macrophages to within 100 μm of the fracture site (F). * = P < 0.05. n ≥ 5 per genotype.

### Patterning of TRAP activity is altered in wnt16^−/−^ zebrafish

TRAP-synthesising osteoclasts are required to resorb damaged bone but must be regulated to prevent osteoporosis. Recombinant WNT16 has been shown to supress osteoclastogenesis and TRAP activity *in vitro* by regulating osteoprotegerin expression in osteoblasts [55]. The uptake of osteoblast-derived extracellular vesicles by immature osteoclasts has been shown to promote osteoclast differentiation in zebrafish scale fractures, demonstrating that intercellular communication between osteoblasts and osteoclasts regulates osteoclastogenesis in response to bone damage [56]. Osteoclasts and macrophages are derived from a common myeloid lineage, with peripheral blood monocytes showing higher osteoclastic potential compared to bone marrow derived monocytes [57]. Moreover, a previous study established that cells expressing the osteoclast marker cathepsin K infiltrate the lepidotrichia fracture site where TRAP is detected by 24 hpi in zebrafish [29]; this coincides with the recruitment of the initial wave of *mpeg1*-expressing cells to the fracture site observed in our model (Figure 4 E-G). Therefore, we investigated whether TRAP activity post-fracture was associated with the recruitment of *mpeg*^*+*^ cells and whether loss of Wnt16 affected levels of TRAP. Fractures were induced in *mpeg1*:mCherry^+^ WT and *wnt16*^*-/-*^ zebrafish and live-imaged prior to amputation of the fin for TRAP staining. The overall levels of osteoclast activity were measured by calculating percentage area of TRAP^+^-stained tissue within 300 μm radius of the fracture site. Osteoclast activity increased rapidly at 24 hpi and remained high before gradually decreasing by 7 dpi (Figure 5 A-B). No significant difference in overall levels of osteoclast activity at the fracture site (TRAP^+^ % area) was detected between WT and *wnt16*^*-/-*^ fractures (Figure 5 B). However, the overall patterning of TRAP staining was altered at 24 hpi and 7 dpi; *wnt16*^*-/-*^ zebrafish displayed a significantly higher number of TRAP^+^ punctae around the fracture, whereas WT fractures tended to display fewer punctae, with continuous, diffuse areas of TRAP^+^ tissue (Figure 5 A & C). Comparable patterning of TRAP^+^ punctae was not observed in uninjured bone from either WT *wnt16* mutants. Interestingly, we observed similarities in the patterning of TRAP^+^ punctae and *mpeg1*^+^ cells, with punctae colocalising with *mpeg1*^*+*^ expression in some regions (Supplementary figure 5). This suggests that *mpeg1*-expressing cells may contribute to bone remodelling and TRAP-synthesis during the early stages of fracture repair.

**Figure 5.**
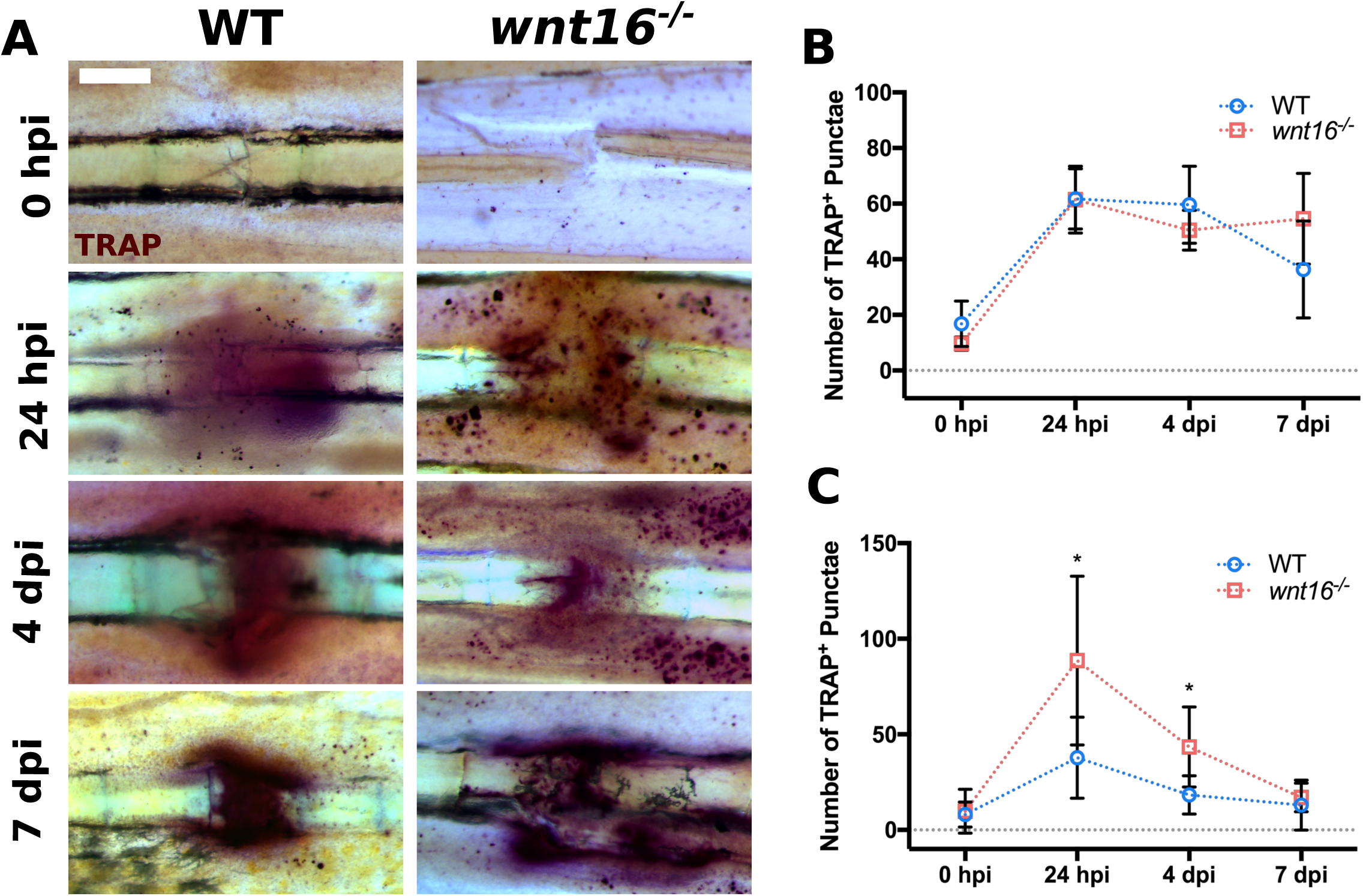
TRAP^+^ punctae accumulate near to fractures in *wnt16*^*-/-*^ zebrafish post-injury. Fins from wild type (WT) and *wnt16*^*-/-*^ zebrafish were amputated at 0 hours post-fracture (hpi), 24 hpi, 4 days post-fracture (dpi) and 7 dpi before undergoing staining to detect the presence of tartrate-resistant acid phosphatase (TRAP). **A:** Representative images of fractures stained for TRAP. Scale bar = 100 *µ*m **B:** Overall coverage of TRAP was measured by calculating the total % area stained within 300 μm of the fracture site. No significant difference in the amount of TRAP^+^ stained area between WT and *wnt16*^*-/-*^ fractures was found. **C:** The number of TRAP^+^ punctae present within 300 μm of the fracture site were quantified and showed a significantly higher number of punctae at 24 hpi and 4 dpi in the fractures of *wnt16*^*-/-*^ zebrafish compared to WT. * = P < 0.05. n ≥ 6 per genotype.

## Discussion

Multiple studies have associated mutations in *WNT16* with osteoporosis and fracture susceptibility phenotypes in humans [15-17], but less was known about the pathophysiological influence of *WNT16* on bone and fracture repair. Moreover, models to study the influence of GWAS-derived fracture-susceptibility candidate genes on bone dynamically *in vivo* were lacking. In this study, we show that loss of Wnt16 in zebrafish leads to variable TMD and the accumulation of bone calluses within lepidotrichia resulting from fractures at an early age. Induction of fractures in caudal fin lepidotrichia and subsequent live imaging showed that Wnt16 is required for optimal fracture healing and the rapid proliferation of osteoblasts post-injury. This coincided with prolonged activation of the canonical Wnt signalling pathway in *wnt16* mutants. Overall, we found that loss of Wnt16 did not impair the development of leukocytes or the responsiveness of neutrophils and macrophages to bone injury but does alter the patterning of TRAP activity at the fracture site.

Disordered activation of the canonical Wnt signalling pathway has been linked to the pathogeneses of many age-related diseases such as cancer, cardiovascular disease, osteoarthritis and osteoporosis [58]. It is thought that WNT16 may antagonise canonical Wnt activity; WNT16 was found to be protective against excessive activation of canonical WNT and severe cartilage degeneration in an induced osteoarthritis murine model [12]. Canonical Wnt signalling culminates in the accumulation of β-catenin in the cell which translocates to the nucleus where it binds to and activates the transcription factors, TCF/LEF (T cell factor/lymphoid-enhancing binding factor). Whilst canonical Wnt signalling is required for osteogenesis, LEF-1 is downregulated in the early stages of fracture repair during soft callus formation [59]. It has been shown that constitutive β-catenin mediated activation of LEF-1 represses the osteoblast transcriptional regulator, Runx-2 and subsequent of expression osteocalcin in osteoblasts [60]. This suggests that delayed callus formation post-fracture in *wnt16*^−/−^ zebrafish is due to overactivation of the canonical Wnt signalling pathway which supresses osteoblast differentiation and bone matrix production.

In contrast to *wnt16* morphant embryos in Clements *et al*., which displayed severe impairment of haematopoiesis [21], we found that loss of Wnt16 had no effect on the overall number of leukocytes detected in larvae during early skeletogenesis, nor did it have an overall effect on the recruitment of neutrophils and macrophages post-fracture. Evidence has shown that gene knockdown using morpholinos can have off-target effects which are hard to control for and show more extreme phenotypes compared to stable mutant lines [61]. Our data demonstrates that targeted, stable loss of Wnt16 via CRISPR-Cas9 mutagenesis does not impair early leukocyte haematopoiesis or the innate immune response to bone injury in adult tissues.

Our data suggests that fracture repair in zebrafish lepidotrichia comprises 3 phases, similarly to mammals [51]. The first is an initial inflammatory phase (∼4-48 hpi) whereby neutrophils and macrophages infiltrate the fracture. This is proceeded by a repair phase (∼3-10 dpi) whereby osteoblasts are activated and synthesise new bone matrix to unionise the fracture with a callus. Ultimately, the bone enters an ongoing remodelling phase (>10 dpi) in which, like humans, the repaired bone remains marked with a calcified callus. Moreover, the biphasic response of macrophage recruitment we observed for the first time in zebrafish is reminiscent of mammalian bone repair, whereby M1-like macrophages are observed during the inflammatory phase and replaced by reparative M2-like macrophages which contribute towards bone matrix synthesis and remodelling of bone [23, 57, 62]. Interestingly, we observed the presence of TRAP^+^ punctae at the fracture site, which coincided with the recruitment of *mpeg1*^+^ macrophages. This suggests that *mpeg1*^+^ cells recruited to the fractures may differentiate into osteoclasts. Indeed, monocytes are known to differentiate into osteoclasts under pro-inflammatory conditions in mammals, whilst Wnt16 has been shown to inhibit the differentiation of bone marrow cells into osteoclasts *in vitro* [22, 55]. Recently, evidence has emerged demonstrating that *mpeg1* expression is not restricted to macrophages in adult zebrafish, as a large proportion of *mpeg1*^*+*^ cells were found to be B cells [63]. The number of TRAP^+^ punctae at the fracture site 24 hpi and 4 dpi was significantly higher in *wnt16* mutants compared to WT. Taken together, this data poses the possibility that *mpeg1* is expressed by other HSC-derived lineages such as osteoclasts, the differentiation of which may be regulated by Wnt16. However, whether distinct sub-populations of macrophages contribute differentially throughout fracture repair in zebrafish requires further investigation.

Though the presence of glycosaminoglycans has been previously reported in the early stages post-injury in a zebrafish fracture model [29], we observed no marked increase in chondrocyte activity throughout fracture repair in the caudal fins of *col2a1*:mCherry zebrafish. This suggests that lepidotrichia bone repair in zebrafish occurs predominantly via intramembranous ossification as opposed to endochondral ossification, unlike mammalian bones, which utilise both mechanisms [64]. Despite this, our study helps to establish zebrafish as a strong, emerging model for studying factors influencing the dynamic behaviour of the multiple cell-types underpinning fracture repair and bone pathologies *in vivo*. By studying the lepidotrichia in the transparent fins of live zebrafish, we were able to visualise bone fragility phenotypes in a novel *wnt16*^*-/-*^ mutant as well as the influence of *wnt16* on bone repair in a dynamic, longitudinal manner. Using this model, we found evidence to suggest that the osteoporosis-associated gene *wnt16* elicits a protective effect against fracture susceptibility and promotes bone repair by buffering levels of canonical Wnt activity to promote optimal osteoblast activity.

## Supplementary materials

Supplementary figures S1-5.

## Materials availability

Mutant *wnt16*^*-/-*^ zebrafish available upon request to the lead contact.

## Data availability

Imaging data will be made available through the university of Bristol’s RDSF server.

## Lead contact

Further information and requests for materials associated with this study should be directed to and will be made available upon reasonable request by the lead contact, Chrissy Hammond.

## Author contributions

Experimental design: LMM, EK, CLH. Conducting experiments: LMM, EK, AV. Image analysis: LMM, AV, SC, EN. Statistical analysis: LMM. Manuscript: LMM, EK, CLH.

## Acknowledgements

We would like to thank Mathew Green and technical staff at the zebrafish aquarium within the University of Bristol’s Animal Scientific Unit for providing animal husbandry and management. We gratefully acknowledge the Wolfson Bioimaging Facility for imaging support.

