## Supplementary figures and images for "Wnt16 Elicits a Protective Effect Against Fractures and Supports Bone Repair in Zebrafish"

### Supplementary Materials S1-5

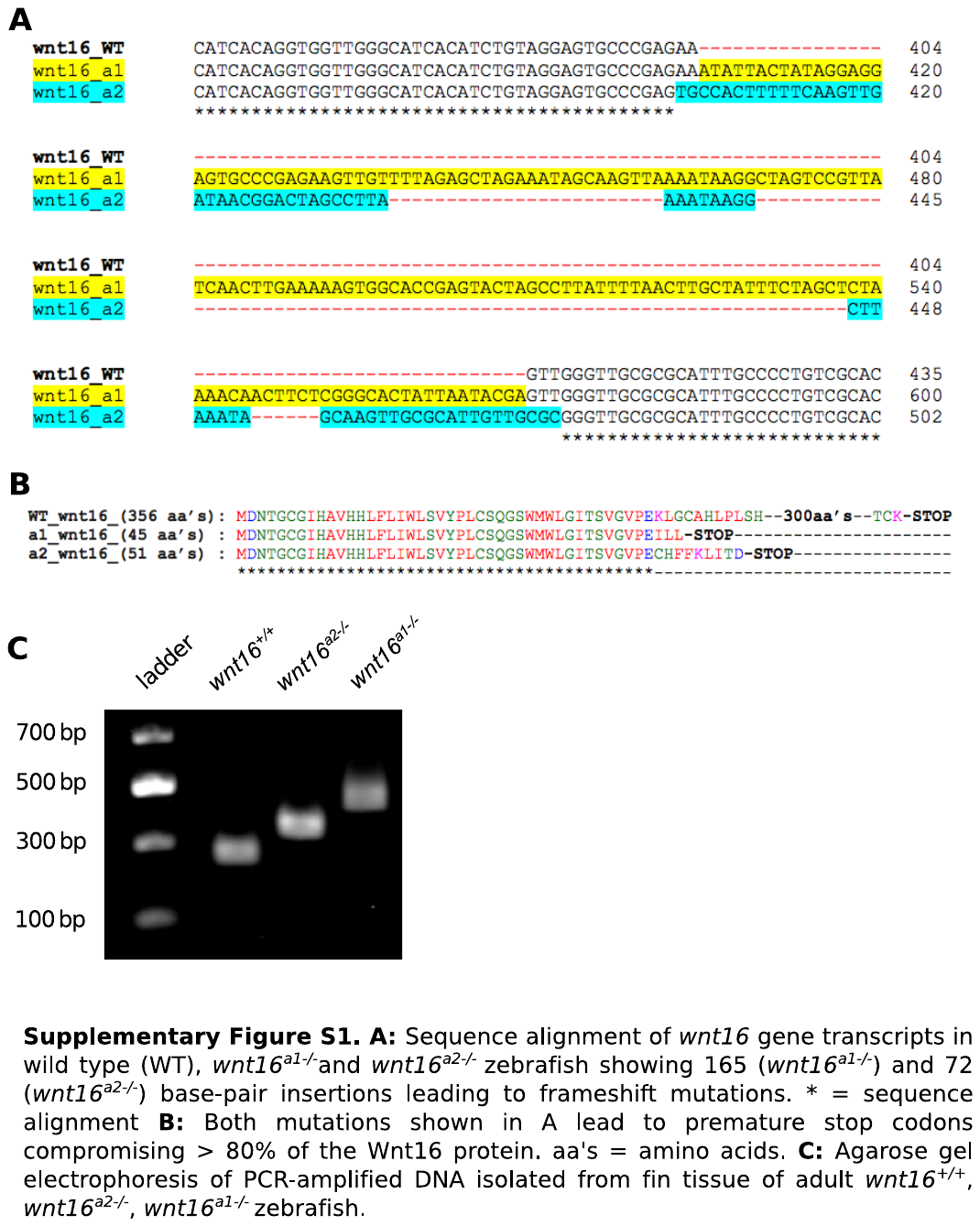

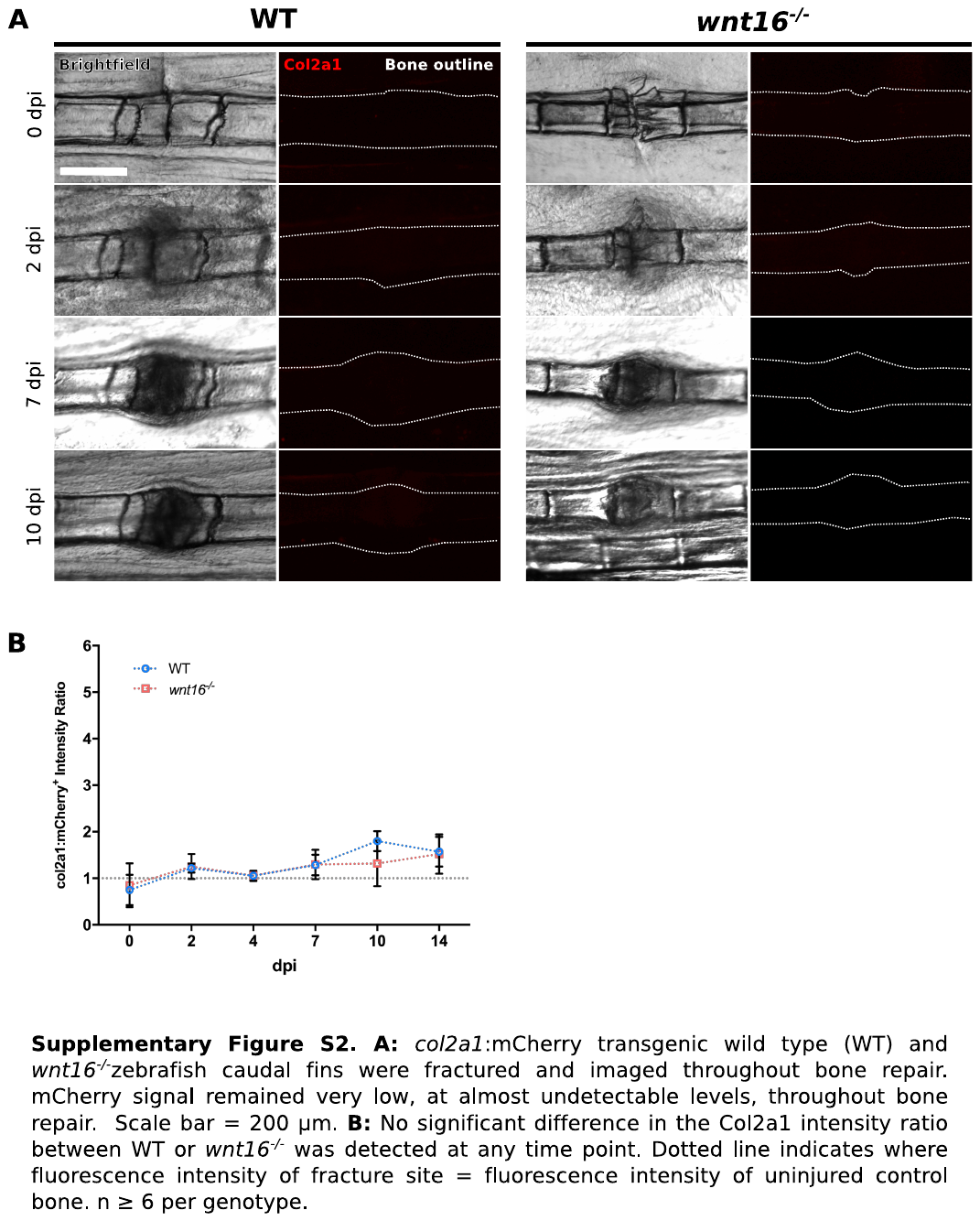

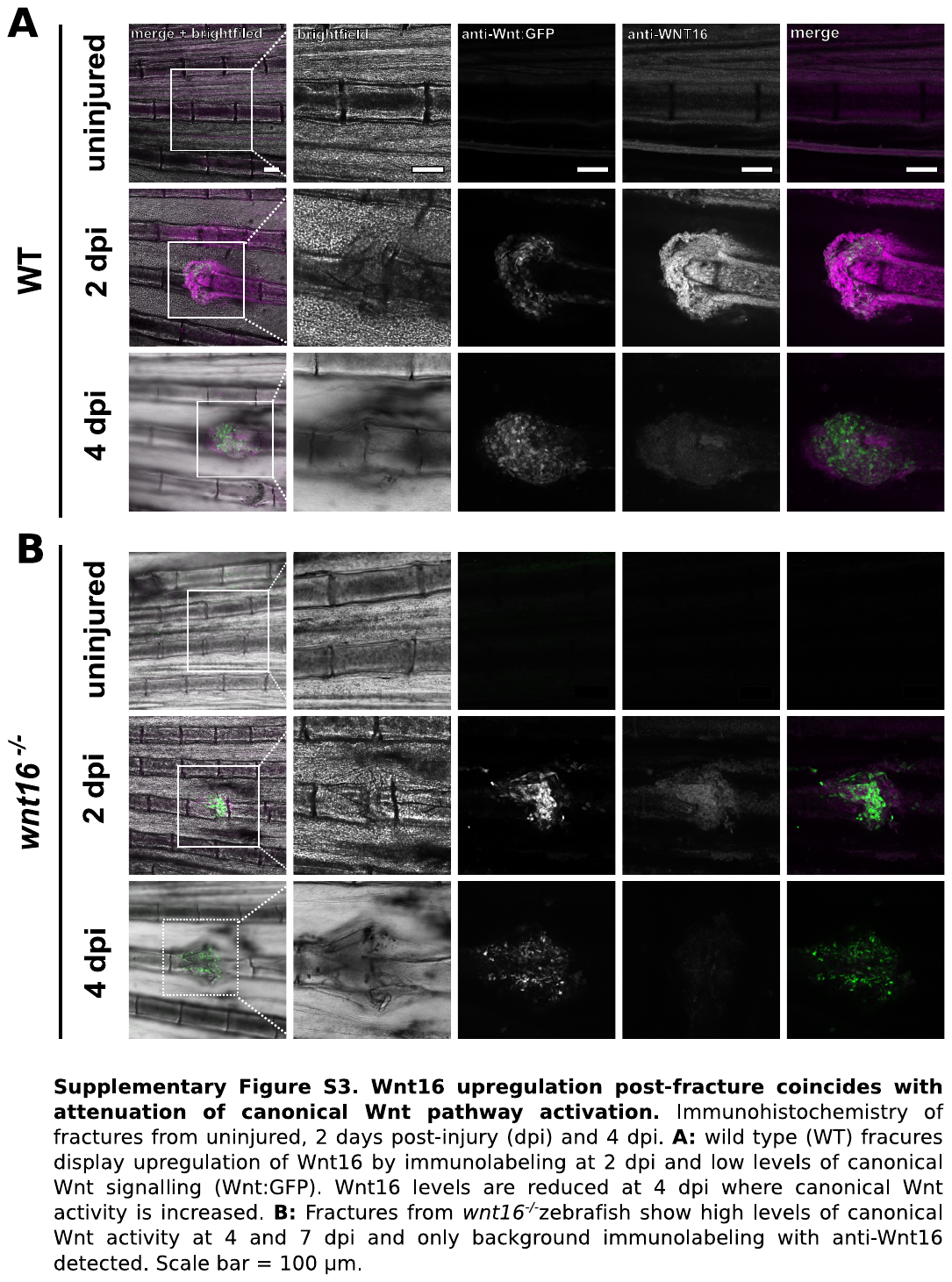

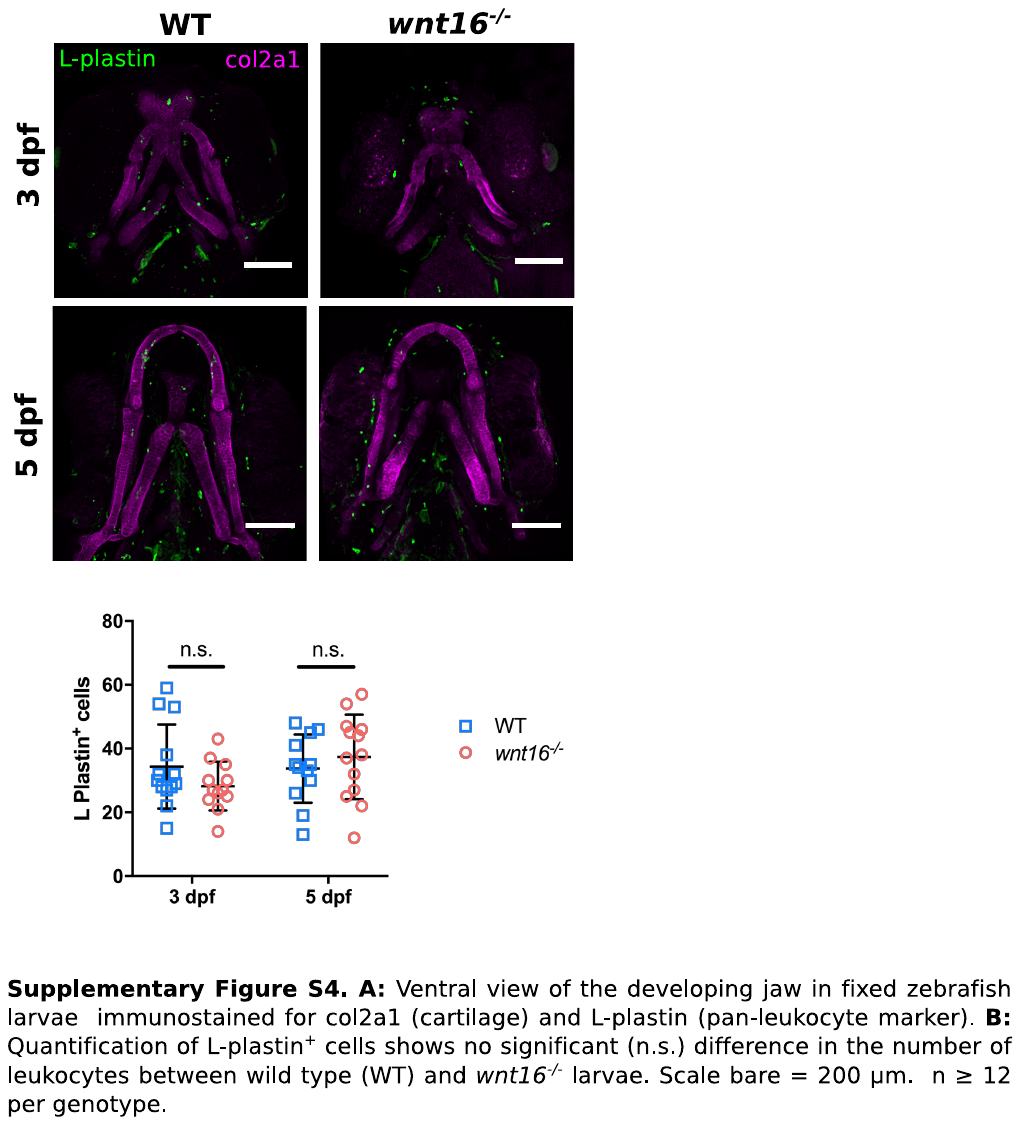

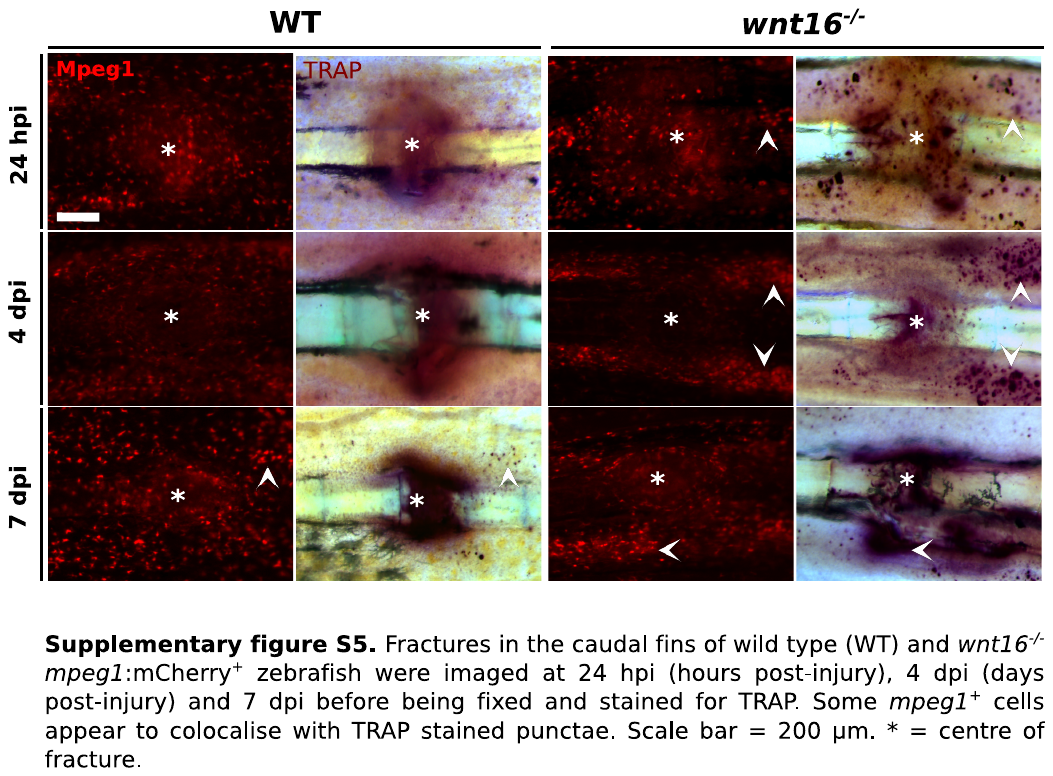
